# Grid cells perform path integration in multiple reference frames during self-motion-based navigation

**DOI:** 10.1101/2023.12.21.572857

**Authors:** Jing-Jie Peng, Beate Throm, Maryam Najafian Jazi, Ting-Yun Yen, Hannah Monyer, Kevin Allen

**Affiliations:** Medical Faculty of Heidelberg University and German Cancer Research Center

**Author notes:** These authors contributed equally (Jing-Jie Peng, Beate Throm). These authors contributed equally (Hannah Monyer, Kevin Allen). Corresponding Author: Dr. Kevin Allen.

## Abstract

With their periodic firing pattern, grid cells are considered a fundamental unit of a neural network performing path integration. The periodic firing patterns of grid cells have been observed mainly during behaviors with little navigational demands, and the firing patterns of grid cells in animals navigating 2D environments using path integration are largely unknown. Here, we recorded the activity of grid cells in mice performing the AutoPI task, a task assessing homing based on path integration. Using artificial deep neural networks to decode the animal’s moment-to-moment movement vectors, we found that grid cells perform path integration over short trajectories and change their reference frames within single trials. More specifically, grid cell modules re-anchor to a task-relevant object via a translation of the grid pattern. The code for movement direction in grid modules drifts as the animal navigates using self-motion cues, and this drift predicts the homing direction of the mouse. These results reveal the computations in grid cell circuits during self-motion-based navigation.

## Introduction

Path integration is a cognitive process whereby an animal estimates its movement from self-motion cues to maintain an up-to-date representation of its position and orientation (Etienne & Jeffery, 2004; McNaughton et al., 1996, 2006; Mittelstaedt & Mittelstaedt, 1980). This process is active whenever an animal moves and contributes to forming new cognitive maps (McNaughton et al., 1996; Savelli & Knierim, 2019). In environments with salient landmarks, animals use a combination of path integration and landmark-based strategies to keep track of their position (Campbell et al., 2018; Jayakumar et al., 2019). When landmarks are absent, the position and orientation of an animal are estimated from path integration alone. In these conditions, the quality of these estimates degrades over time due to error accumulation (Chen et al., 2016; Pérez-Escobar et al., 2016; Stangl et al., 2020), making navigation using path integration alone more suited for short navigation journeys.

Lesion studies have shown that the medial entorhinal cortex, the hippocampus, and the retrosplenial cortex all contribute to the ability of an animal to navigate using path integration (Elduayen & Save, 2014; Maaswinkel et al., 1999; Parron & Save, 2004; Van Cauter et al., 2013). The activity patterns of neurons within these brain regions that support navigation using path integration in 2D environments have yet to be established. The functional cell type most often linked to path integration is, without a doubt, the grid cells located in the medial entorhinal cortex (Burak & Fiete, 2009; Clark & Nolan, 2023; Gil et al., 2018; Hafting et al., 2005; McNaughton et al., 2006; Sorscher et al., 2023; Ying et al., 2022). Grid cells have several firing fields organized as one continuous grid of equilateral triangles. It has been proposed that these cells provide a global coordinate system in which the current location and orientation of an animal are updated via self-motion cues (Banino et al., 2018; Bush et al., 2015; Erdem & Hasselmo, 2012; McNaughton et al., 2006; Stemmler et al., 2015). However, it is still unclear whether the continuous grid firing pattern of grid cells is present in animals navigating purely by path integration, as this grid pattern has been observed mainly in animals performing random foraging tasks in open-field environments or navigation tasks with salient landmarks (Boccara et al., 2019; Butler et al., 2019).

Here, we set out to characterize the firing patterns of grid cells in mice navigating using path integration and tested whether grid cell activity predicted homing behavior. We took advantage of the newly established AutoPI homing task, which allows large-scale electrophysiological recordings of spatially selective neurons in mice navigating a 2D environment utilizing path integration. Moreover, we developed a deep-learning decoding framework to monitor the path integration process in grid cell modules with high temporal resolution and showed that grid map orientation predicts homing behavior.

### Absence of stable grid patterns in mice navigating using self-motion information

We recorded extracellular action potentials in the MEC of 17 mice as they performed random foraging trials and the AutoPI task (Najafian Jazi et al., 2023) (**Fig. 1a top**). On each trial of the AutoPI task, the mouse left a home base to search for a lever on the arena, pressed the lever, and returned to the home base to collect its food reward (**Fig. 1a bottom**). The mouse performed AutoPI trials under normal lighting conditions and in darkness (light and dark trials). We analyzed 180 recording sessions in which mice performed between 38 and 172 trials per recording session. Mice had longer search paths to the lever during dark than light trials (**Fig. b**). The ability of mice to return directly to the home base after pressing the lever was quantified with the homing error at the periphery (**Extended Data** Fig. 1a). Error at the periphery was larger during dark than light trials (**Fig. 1c**, **Extended Data** Fig. 1b-c), but it remained above the chance level during dark trials (chance level = π/2).

**Fig. 1.**
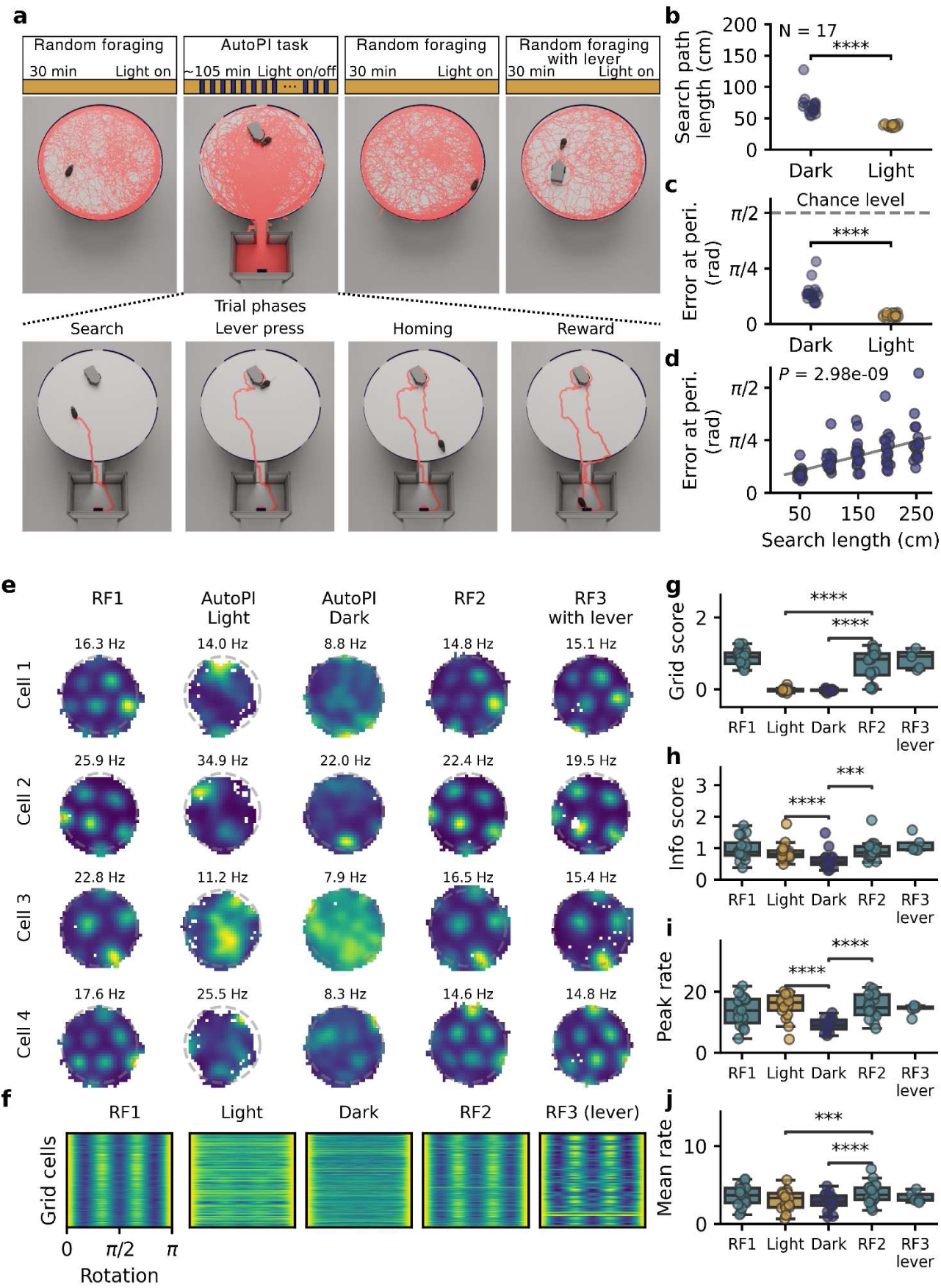
| Reduced grid cell periodicity in mice performing a homing task. **a**, Top: Recording protocol. During each recording session, mice performed a random foraging task before and after the AutoPI task. On the AutoPI task, mice alternated between trials with and without visible light (light and dark trials). In the last recording session of each mouse, mice performed an additional random foraging trial in which the lever box was placed on the arena. The red lines on the arena and floor of the home base are examples of the path of a mouse during a recording session. Bottom: The four phases of a trial on the AutoPI task. The mouse leaves the home base and searches for the lever box on the arena. Once at the lever box, the lever is pressed, triggering food reward delivery in the food magazine of the home base. The mouse returns to the home base (homing) to access the food reward. **b**, Search path length during dark and light trials (Dark-Light difference: N = 17 mice, Wilcoxon signed-rank test, statistic = 0.0, *P* = 1.53 x 10^-5^). **c**, Homing error at the periphery during dark and light trials (N = 17 mice, Wilcoxon signed-rank test, statistic = 0.0, *P* = 1.53 x 10^-5^). **d**, Homing error as a function of search path length during dark trials (N = 17 mice, Friedman test, statistic = 45.6, *P* = 2.98 x 10^-9^). **e**, Firing rate maps of four grid cells during the different tasks (random foraging (RF) and AutoPi task). RF3: random foraging task with the lever on the arena. **f**, Grid cell periodicity matrix in different tasks. Each row of the matrix represents the data from one grid cell. Grid periodicity is reflected by the peaks for the rotations of 0, π/3, and 2*π/3 rad. **g**, Grid scores for grid cells during RF trials and light and dark trials of the AutoPI task (N = 17 mice except for RF3-lever where N = 5 mice, Wilcoxon signed-rank test, Light-RF2 difference: statistic = 0.0, *P* = 1.53 x 10^-5^; Dark-RF2 difference: statistic = 0.0, *P* = 1.53 x 10^-5^). **h**, Information scores of grid cells (Light-RF2: statistic = 45.0, *P* = 0.15; Dark-RF2: statistic = 4.0, *P* = 1.07 x 10^-4^; Light-Dark: statistic = 0.0, *P* = 1.53 x 10^-5^). **i**, Peak firing rate in the firing rate maps of grid cells (Light-RF2: statistic = 62.0, *P* = 0.52; Dark-RF2: statistic = 1.0, *P* = 3.05 x 10^-5^; Light-Dark: statistic = 1.0, *P* = 3.05 x 10^-5^). **j**, Mean firing rate of grid cells (Light-RF2: statistic = 8.0, *P* = 3.81 x 10^-4^; Dark-RF2: statistic = 1.0, *P* = 3.05 x 10^-5^; Light-Dark: statistic = 40.0, *P* = 0.089). \*\*\*\**P* < 0.0001, \*\*\**P* < 0.001.

One signature of behaviors supported by path integration is that navigation error increases with the running path length (Stangl et al., 2020). We tested if this was the case during the AutoPI task. Error at the periphery grew as a function of search path length (**Fig. 1d**, **Extended Data** Fig. 1d-e). This correlation between search path length and homing error was observed independently of how lateral the position of the lever was on the arena (**Extended Data** Fig. 1f, **Extended Data** Fig. 2). These results support the hypothesis that mice used path integration to return to the home base during dark trials.

We recorded 5746 neurons, of which 931 were classified as grid cells based on their significant grid scores during the first random foraging trial (**Fig. 1e, Extended Data** Fig. 3, **Extended Data File 1**). Grid cells showed spatial periodicity during both random foraging trials but, surprisingly, not during the AutoPI task (**Fig. 1e-g**). Grid scores were lower during the light and dark trials of the AutoPI task compared to those observed during random foraging trials (**Fig. 1g**). The decrease in grid periodicity observed during the AutoPI task was not simply the result of the lever being present on the arena. Indeed, when mice performed a random foraging trial in which the lever was on the arena (RF3-lever), grid cells displayed strong periodicity (**Fig. 1e-g**). The information scores and peak firing rates of grid cells were lower during dark trials than light trials of the AutoPI task (**Fig. 1h-i**).

### Moment-to-moment decoding of movement path from grid cell activity

The reduction in grid periodicity during the AutoPI task questions the assumption that the grid cell network performs path integration during the homing task. We tested this directly by developing a computational framework to predict the mouse’s ongoing movement vectors based on grid cell activity (**Fig. 2**). We took advantage of the fact that the activity of grid cells spans the surface of a torus and that the activity manifold is preserved across behaviors (Gardner et al., 2022) (**Extended Data** Fig. 4). We identified simultaneously recorded grid cells from the same module (i.e., with similar orientation and spacing). The first random foraging trial was used to establish the orientation and period of the grid pattern and transform the cartesian position data (x,y) into coordinates in toroidal space (v_0_,v_1_)(**Fig. 2a**). We trained a recurrent neural network to predict the animal’s position in toroidal space based on the instantaneous firing rate of grid cells (**Fig. 2b**). To predict the movement vectors of the mouse, we calculated the change in the predicted toroidal position over time, resulting in a series of movement vectors in toroidal space. These movement vectors were then transformed from toroidal space to cartesian space. The cumulative sum of the cartesian movement vectors represented the predicted movement path (**Fig. 2b, right**). If a grid cell module performs path integration, the real and predicted movement paths resemble each other.

**Fig. 2.**
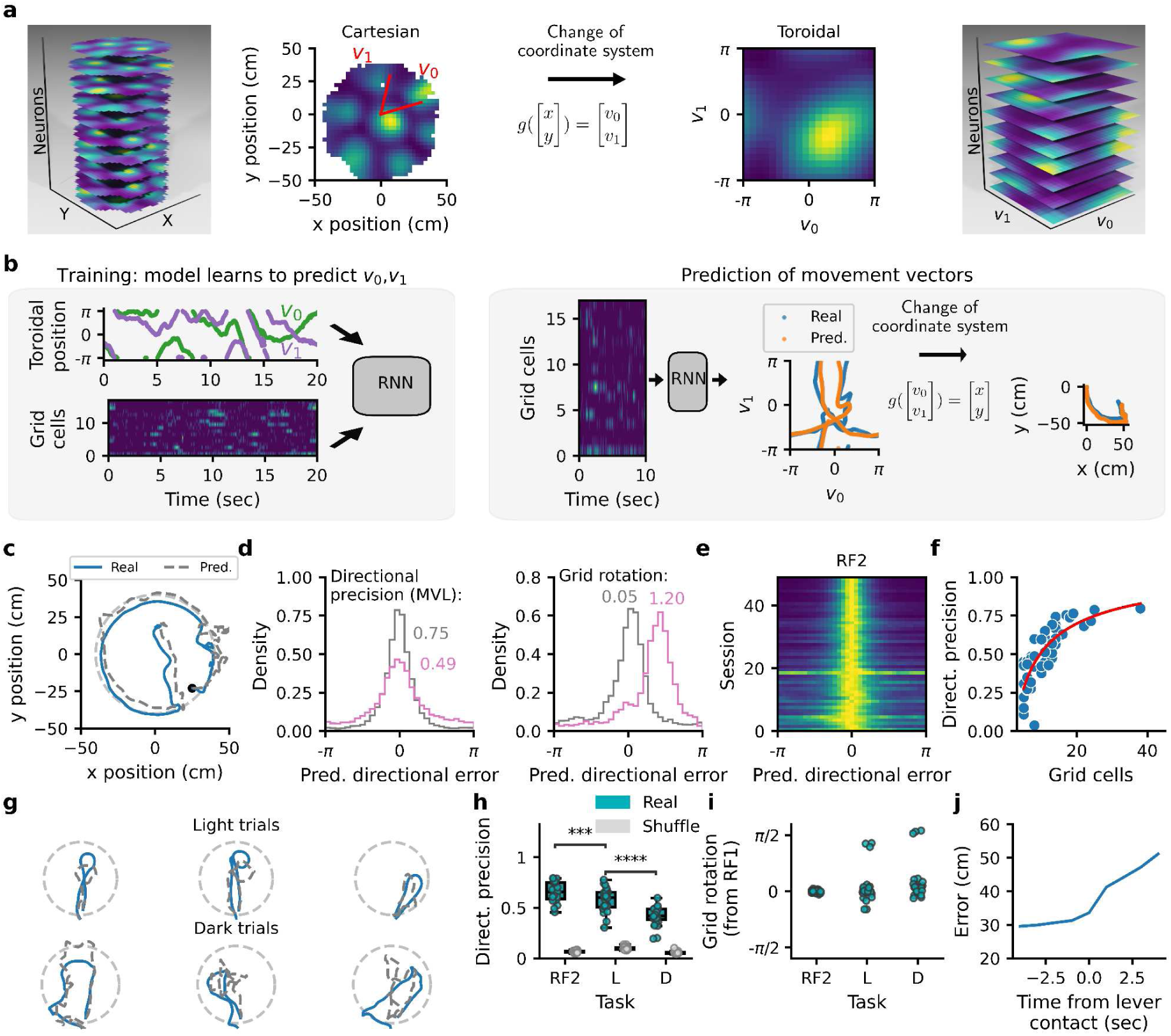
| Prediction of the animal’s movement vectors from grid cell activity. **a**, Transformation of cartesian position data to a toroidal coordinate system. Left: Stack of grid cell firing rate maps from random foraging. Grid orientation and period were estimated to define the two main grid axes (*v0* and *v1*). Right: firing rate maps of the same grid cells in toroidal space. **b**, Left: A recurrent neural network (RNN) was trained to predict the animal’s position in toroidal space (v0 and v1) from the instantaneous firing rate of simultaneously recorded grid cells. The data from the first random foraging trial were used for training. Right: The trained RNN predicted the animal’s position in toroidal space. Toroidal movement vectors were calculated from the change in toroidal position over time. The toroidal vectors were transformed into cartesian vectors and were summed to reconstruct movement paths. **c**, Example of real (blue) and predicted (gray) paths during the second random foraging trial. 25 seconds are shown. **d**, Two measures derived from the distribution of predicted directional error. The data from two sessions are shown (gray and pink). The mean vector length (MVL) of these distributions reflects the directional precision of the model (left), whereas their circular means represent the rotation of the grid representation relative to the first random foraging trial (right). **e**. Distribution of predicted directional error for 49 recording sessions during the second random foraging trial (RF2). **f**, Directional precision as a function of the number of grid cells in the model (Pearson correlation, N = 49 sessions, r = 0.736, *P* = 1.67 x 10^-9^). The data were fitted with a logarithmic equation of the second degree (red curve). **g**, Examples of real (blue) and predicted (gray) paths during the AutoPI task. **h**, Directional precision during the second random foraging (RF2) and light (L) and dark (D) trials on the AutoPI task compared to shuffled data (N = 24 sessions, Wilcoxon signed-rank test, RF2-L: statistic = 35.0, *P* = 0.00049, L-D: statistic = 3.0, *P* = 5.96 x 10^-7^). **i**. Rotation of the grid representation during the second random foraging (RF2) and light (L) and dark (D) trials on the AutoPI task. **j**, Cumulative error in position prediction as a function of time from finding the lever during dark trials. \*\*\*\**P* < 0.0001, \*\*\**P* < 0.001.

**Fig. 2c** shows the similarity between real and predicted paths during random foraging. To quantify the quality of the path prediction, we calculated the angle between the real and predicted movement directions, referred to as predicted directional error. From the distribution of predicted directed error, we extracted two principal measurements: directional precision and grid rotation (**Fig. 2d**). The directional precision was defined as the mean vector length of the predicted direction error distribution, whereas the grid rotation relative to the first random foraging trial was obtained by calculating the circular mean of the predicted directional error distribution. There was no rotation of the grid pattern between the first and second random foraging trials (**Fig. 2e**). The directional precision of the model increased with the number of grid cells participating in the prediction (**Fig. 2f**). For recording sessions with at least 15 grid cells (N = 9), the directional precision when running speed exceeded 10 cm/sec had a median of 0.76, a degree of directional selectivity comparable to sharply tuned head-direction cells (Yoder & Taube, 2009).

We then applied the path reconstruction analysis to the AutoPI task (**Fig. 2g**) for sessions where the directional precision during the second random foraging trial was above 0.5. Directional precision decreased from random foraging to light trials of the AutoPI task and from light trials of the AutoPI task to dark trials (**Fig. 2h**). The directional precision during dark trials of the AutoPI task was higher than chance levels for all sessions in which path reconstruction was attempted (24 out of 24 sessions), demonstrating that grid-cell modules performed path integration. The orientation of the grid representation was maintained from random foraging to the AutoPI task in all but one mouse (**Fig. 2i**). During dark trials of the AutoPI task, the error accumulation rate in the predicted integrated path increased once the mouse reached the (**Fig. 2j**).

### Re-anchoring of grid cells during navigation

During the homing task, we observed decreased path prediction accuracy after the mouse reached the lever (**Fig. 2j**). We hypothesized that this could be caused by a re-anchoring of the grid pattern to the randomly located lever. We first characterized the firing fields of grid cells when the mouse ran around the lever. We found that many grid cells fired when the mouse was located in a specific direction relative to the center of the lever (**Fig. 3a**). Importantly, these fields were present in most trials even though the position of the lever changed across trials. Comparing the firing rate histograms of grid cells around the lever during light and dark trials (**Fig. 3a, second row**), we found that the median within-cell correlation between the histograms was low (n = 512 grid cells, median r = 0.048), indicating that the firing fields of grid cells often had different locations between light and dark trials. The directional selectivity of grid cells around the lever during light or dark trials was significantly higher than chance levels (**Fig. 3b**). These lever-anchored grid fields suggest that grid cells could be influenced by two reference frames on the AutoPI task: a room reference frame and a lever-centered reference frame. To assess their respective impact, we calculated firing rate maps in both reference frames (**Extended Data** Fig. 5a). Surprisingly, during dark trials, the stability of firing rate maps was higher in the lever-centered reference frame than in the arena reference frame (**Fig. 3c**).

**Fig. 3.**
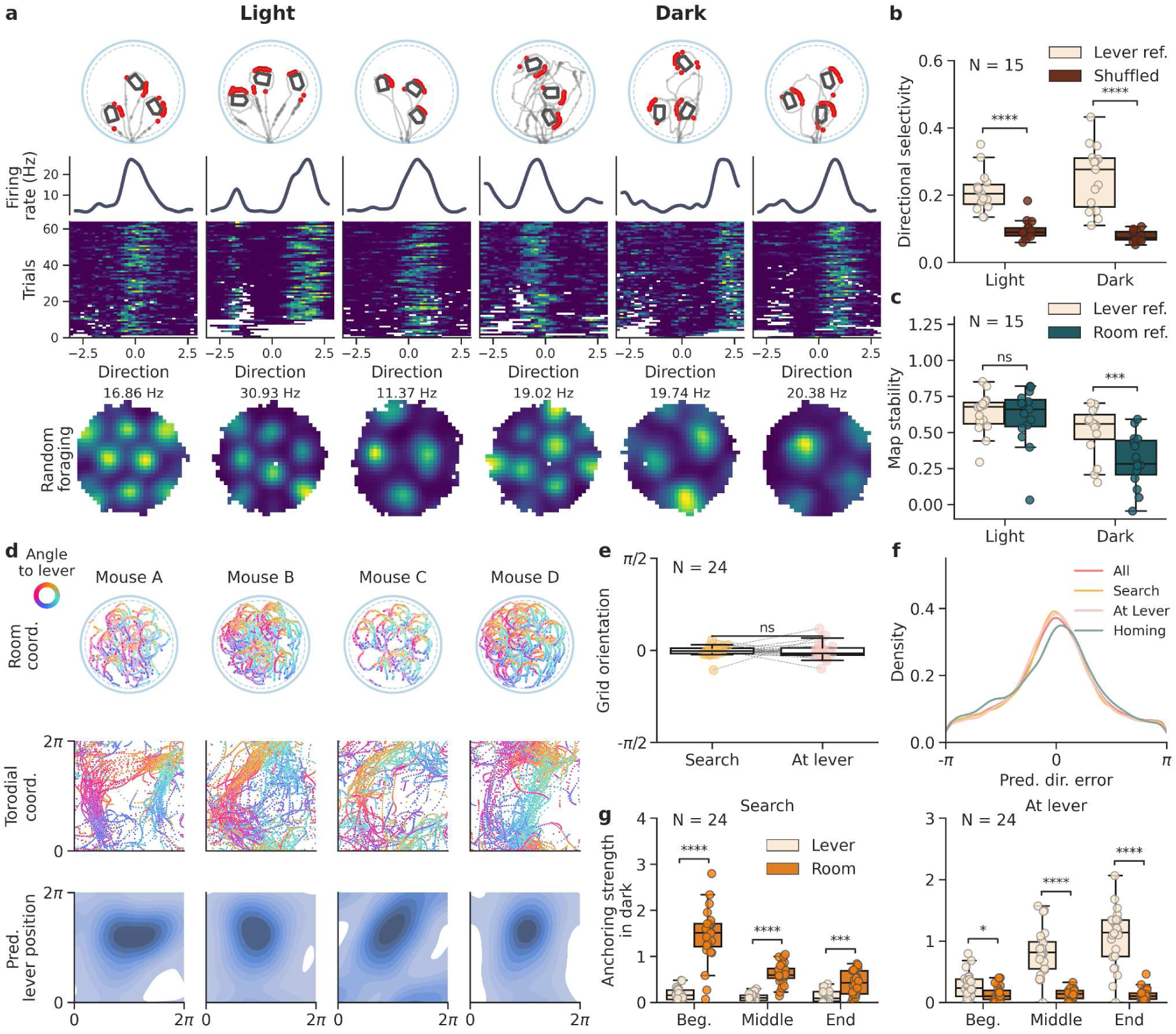
| Re-anchoring of grid cell modules to the lever location. **a**, Examples of six grid cells with firing fields near the lever location in light and dark trials. First row: Spikes (red) on the running path (gray) during three trials. Only spikes emitted within 18 cm of the lever are shown. Second row: Firing rate as a function of the direction of the animal relative to the lever center. Third row: Firing rate as a function of direction around the lever on different trials. Each row represents a trial in which the mouse pressed the lever. Fourth row: Firing rate maps during random foraging. **b**, Directional selectivity (MVL) of neurons in the lever-centered reference frame against shuffling in light and dark trials (N = 15 mice with grid cells with lever-anchored fields, Wilcoxon sign-rank test, light: *P* = 6.104 x 10^-5^, dark: *P* = 6.104 x 10^-4^). **c,** Map stability of grid cells with lever-anchored firing fields. Maps were calculated in the room reference frame or the lever reference frame (N = 15 mice with grid cells with lever-anchored fields, Wilcoxon sign-rank test, light: *P* = 9.780 x 10^-1^, dark: *P* = 8.545 x 10^-4^). **d,** Anchoring of grid cell modules to the lever position during dark trials. Top: Examples of the running paths of the mouse around the lever during recording sessions from four different mice. The mouse’s path is color-coded to reflect the direction of the mouse around the lever. Middle: Predicted position of the mouse in toroidal space when the mouse is around the lever. The color code represents the direction of the mouse relative to the lever. Bottom: Kernel density estimate of the predicted lever position in toroidal space when the mouse is near the lever. **e,** Grid orientation relative to random foraging during search behavior and when the mouse was near the lever. The data are from dark trials. **f,** Distribution of predicted directional error for all dark trials or during three separate phases (search, at lever, and homing) of the dark trials. **g,** Anchoring strength to the lever or the room reference frame during dark trials. Data during the search behavior and when the mouse was at the lever are shown separately and split evenly into the beginning, middle, and end (N = 24 sessions, Wilcoxon sign-rank test, Search, Beg.: *P* = 5.515 x 10^-8^, Middle: *P* = 1.178 x 10^-8^, End: *P* = 1.532 x 10^-4^; At lever, Beg.: *P* = 1.365 x 10^-2^, Middle: *P* = 4.611 x 10^-8^, End: *P* = 7.371 x 10^-8^). \*\*\*\**P* < 0.0001, \*\*\**P* < 0.001, * 0.01 < *P* < 0.05, ns 0.05 < *P*.

We tested the hypothesis that grid cell modules anchor to the lever position using the RNN predicting the animal position in toroidal space (**Fig. 2b**). At the grid module level, anchoring to an object implies that this object has a stable location in toroidal space, irrespective of the location of the object on the arena. **Fig. 3d** shows the animal’s position as it ran around the lever in cartesian space and the predicted position in toroidal space (**Fig. 3d top and middle**). In several recording sessions, the predicted paths in toroidal space, when the animal ran around the lever formed a ring. This ring was particularly evident during dark trials. To estimate the position of the lever in toroidal space, we obtained the cartesian vector from the animal’s position to the lever and added the equivalent toroidal vector to the predicted toroidal position of the mouse (**Extended Fig. 5b**). When the mouse was near the lever, the lever position in toroidal space clustered at a specific location (**Fig. 3d bottom**). We fitted a bivariate von Mises distribution to the lever positions in toroidal space and used the two kappa parameters to measure grid anchoring strength. During dark trials, the grid anchoring to the lever was statistically significant in 23 out of 24 sessions (**Extended Fig. 5d**). The peaks of the predicted directional error distribution were centered on 0, indicating that there was no systematic rotation of the grid pattern when the mouse was around the lever (**Fig. 3e-f**). This demonstrates that the re-anchoring to the lever occurred via a translation of the grid pattern.

Using the same method to assess grid anchoring strength, we assessed how grid anchoring evolved during single runs on the arena (**Fig. 3g**). Grid modules were firmly anchored to the room reference frame at the beginning of the search path, but the dominant anchoring transitioned to the lever position after the mouse reached the lever. After departing from the lever, the degree of anchoring to the lever decreased, particularly in the later stages of the homing behavior (**Extended Data** Fig. 5f).

### Grid orientation drift increases with running path length

Although we found that the mean orientation of the grid pattern did not change from random foraging to the AutoPI task (**Fig. 2i and 3e**), we hypothesized that, in individual trials in darkness, the orientation of the grid pattern drifted from its original orientation. We tested whether this directional drift increased as the animal searched for the lever in darkness. We compared the predicted directional error of the model (as in **Fig. 2d**) during the beginning and end of the search paths of dark trials (**Fig. 4a**). The directional precision of the model was higher at the beginning of the search compared to the end (**Fig. 4a,b**), suggesting that the orientation of the grid pattern drifts during the search path. We also observed a progressive increase in the absolute predicted directional error as a function of search duration (**Fig. 4c**). The directional precision when the mouse was at the lever was lower after long search paths than shorter ones (**Fig. 4d**). We also quantified the directional drift on single trials as the animal ran around the lever in darkness and found that the directional drift was higher after long than short search paths. (**Fig. 4e**). These results are consistent with the idea that grid cell orientation during dark trials is updated by an angular path integrator that accumulates error over time or distance run. Importantly, despite re-anchoring events, the accumulated error by the path integrator during the search is preserved after the change of reference frames.

**Fig. 4.**
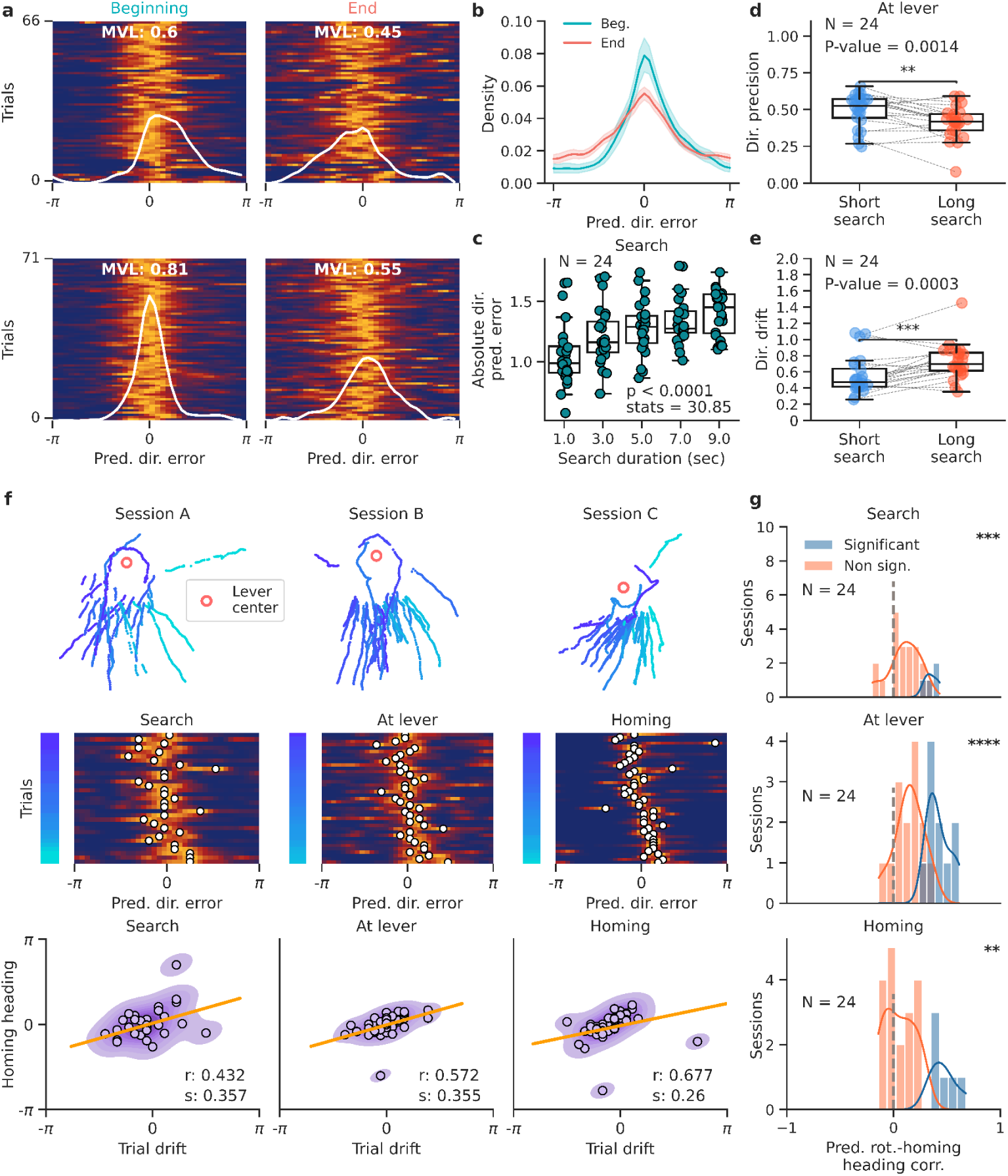
| Grid cell orientation predicts homing behavior. **a**, Trial matrices of the predicted directional error during long search paths’ beginning (left) and end (right) of dark trials. Two recording sessions are shown, one per row. The white line shows the mean predicted directional error across trials. The mean vector length (MVL) of the distribution is shown. **b,** Probability density of the predicted directional error during the beginning and end of long search paths (N = 24 sessions, Kolmogorov-Smirnov Test, statistic = 0.265, P = 4.359 x 10^-27^). **c,** Absolute predicted directional error as a function of search duration during dark trials (N = 24 sessions, Friedman test, statistic = 30.85, P = 9.141 x 10^-7^). **d,** Directional precision when the mouse was at the lever after short or long search paths (N = 24 sessions, Wilcoxon sign-rank test, *P* = 1.400 x 10^-3^). **e**, Directional drift when the mouse was at the lever after short and long search paths (N = 24 sessions, Wilcoxon sign-rank test, *P* = 2.781 x 10^-4^). **f,** Examples of correlation between the predicted directional error and homing direction. First row: lever-centered homing paths of the animal in three sessions. Paths are color-coded by the mouse’s median heading direction while homing. Second row: trial matrix of the predicted directional error on single trials during the search (left), when the mouse was at the lever (middle), or during homing (right). The trials were sorted by the homing heading of the mouse, as indicated by the color code on the left of each matrix. A white circle indicates the peak in the predicted directional error distribution. Third row: predicted directional error against homing heading error. Points represent individual dark trials. The predicted directional error was calculated when the mouse was at the lever (middle), or during homing (right). A regression line (orange), the circular correlation coefficient (r), and the slope (s) are shown. **g,** Distribution of circular correlation coefficients between the predicted directional error and homing heading during the search path (top, stats = 38.0, P = 7.423 x 10^-4^), when the mouse was at the lever (middle, stats = 7.0, P = 2.265 x 10^-6^), and during the homing path (bottom, stats = 53.0, P = 4.335 x 10^-3^). \*\*\*\**P* < 0.0001, \*\*\**P* < 0.001, ** 0.001 < *P* < 0.01

### Grid orientation drift predicts homing behavior

We investigated whether the grid orientation drift that accumulated during the search and as the animal ran near the lever predicted the direction of the mouse’s homing behavior. We calculated the predicted directional error of the model separately during the search, while the mouse was at the lever, and during homing (**Fig. 4f**). On each dark trial, we identified the peak in the predicted direction error distribution, which we referred to as trial drift. We found that the trial drift was correlated with the homing heading direction (**Fig 4f**). Notably, the trial drift of grid modules during the search, when the animal was around the lever, and during homing behavior was positively correlated with the animal’s homing heading direction (**Fig. 4f**). The distribution of r values for the *trial drift/homing direction* correlations was significantly shifted towards positive values (**Fig. 4g**), demonstrating that the trial drift of the grid map during the search path, at the lever, or during the homing path all predicted homing direction.

## Discussion

Our findings demonstrate that grid cells do not maintain a stable grid representation during a path integration task. Instead, we observed a transition from a room reference frame to a local reference frame centered on the location of the lever. This re-anchoring of the grid map involved a translation of the grid pattern with no change in grid orientation. Interestingly, a selective grid phase change without changes in grid orientation has already been reported after changing non-metric cues or the shape of an environment (Fyhn et al., 2007; Marozzi et al., 2015). This indicates that the grid phase can anchor to multiple local reference frames while grid orientation remains constant. Here, we demonstrate that after short exploratory journeys in which position is estimated mainly from self-motion cues, the grid phase is re-anchored whenever a known landmark is encountered, even if this landmark does not have a stable position. This anchoring of the grid maps to the lever could explain why a quarter of CA1 pyramidal cells, located one or a few synapses downstream from grid cells (Zhang et al., 2013), also have lever-anchored firing fields during this same behavioral task (Najafian Jazi et al., 2023). We predict that grid cells change their anchoring point in many navigational tasks where place cells have been shown to change reference frames (Gothard, Skaggs, & McNaughton, 1996; Gothard, Skaggs, Moore, et al., 1996; Moore et al., 2021).

Although stable grid patterns were not observed during the AutoPI task, we could predict movement direction with a significant level of accuracy, demonstrating that grid cell modules performed path integration while the mouse ran on the task. The lack of a stable grid pattern was caused by re-anchoring the grid phase when the animal reached the lever and noise accumulation in the estimation of orientation and position when navigating using self-motion cues (Chen et al., 2016; Hardcastle et al., 2015; Pérez-Escobar et al., 2016).

The computations performed by grid cells during memory or navigation tasks have been notoriously difficult to characterize because standard measurements of grid cell activity require recording periods of several minutes, whereas behaviorally relevant events often last only a few seconds. With our computational framework to monitor grid cell activity in toroidal space, we could infer the grid orientation and grid phase anchoring with a sub-second resolution. This opens up the possibility of characterizing the properties of grid cells at the module level during most standard navigation and memory paradigms by removing the need for long behavioral epochs with extended spatial coverage. This approach, and similar module-level analysis (Gardner et al., 2022; Hermansen et al., 2022; Wen et al., 2023), will contribute to understanding grid cell functions in a broader spectrum of behaviors.

## Methods

### Apparatus for the AutoPI task

The behavioral apparatus consisted of a home base and a circular arena, both elevated 45 cm above the floor. The home base was a rectangular box (20 × 30 × 30.5 cm) with a front-facing sliding door and a food magazine attached to its back wall. The food magazine was equipped with an infrared beam to detect the presence of the mouse at the magazine. Food rewards (AIN-76A Rodent tablets 5 mg, TestDiet) were delivered into the magazine from a pellet dispenser attached to the external side of the home-base outside wall. The sliding door of the home base could be lowered to give access to a bridge (10 x 10 cm) leading to the arena.

The circular arena (diameter: 80 cm) was mounted on a tapered roller bearing to allow clockwise and anti-clockwise rotations. Eight 1.6 cm high wall inserts were attached to the edge of the arena. The space between adjacent inserts was 10 cm, creating eight evenly spaced openings that potentially led to the bridge. One of the eight openings always faced the home base bridge when the mouse was on the arena. The arena rotation, movement of the door, and delivery of food rewards were operated via Arduino Uno microcontrollers, digital stepper motor drivers (Stepperonline, DM542T), and N17 stepper motors. Two cameras (above the home base and the arena) connected to a microcomputer (Jetson Xavier Nx) were used to monitor the animal and lever during the task. The videos were captured at 30 Hz (640 × 480 pixels), analyzed online, and stored for further offline processing.

A lever (13 x 10 mm) extended from one wall of an autonomous lever box (116 × 82 × 80 mm). The lever box contained two servo motors, an Arduino Nano connected to a radio frequency module (NRF24L01), and a 2000mAh lithium battery. Pressing the lever broke an infrared beam inside the lever box. Lever press timestamps were transmitted via the radio module to a microcomputer controlling the task (Jetson Xavier Nx). The same radio module was used to instruct the lever box to move to a different location on the arena. The location and orientation of the lever box were obtained using DeepLabCut (https://github.com/DeepLabCut/DeepLabCut). When moving the lever between trials, the intended future location and orientation were randomly selected within a circle centered on the arena center with a radius of 75% of that of the arena. The Arduino Nano inside the lever box was instructed to move along the vector ending at the new locations. The new position and orientation were confirmed via DeepLabCut and corrected if needed.

Trials in the AutoPI task were performed with and without visible light and were referred to as light and dark trials, respectively. During light trials, 2 LED stripes (30cm, white) above the arena were the only visible light source. An Arduino equipped with a relay module turned them on and off. During dark trials, the only light sources were infrared light LED stripes and lamps (850 µm) positioned above the arena and the home base. A white noise (65 dB) generated from a speaker above the arena was played during the task to mask uncontrolled auditory cues.

A Python script running on a microcomputer (Nvidia Jetson Nx) controlled the logic of the task. Modular computer programs were responsible for controlling the different elements of the task (door movement, arena rotation, visible light, lever position, and food delivery). The Robot Operating System (ROS, https://www.ros.org) established communication between these nodes. Task-related events (e.g., lever press, food delivery, door movement, arena rotation, light switch) and their respective time stamps were saved into a log file.

### Subjects

The subjects were 17 5-9 months old C57BL/6 male mice. The mice were kept on a 12-hour light/12-hour dark cycle, with all experiments performed during the light period. The mice were single-housed in 26 x 20 x 14 cm plexiglass cages with food and water *ad libitum*. The cage contained 2-3 cm of sawdust and was enriched with paper tissues. The experimenters handled the mice for 5-10 minutes, once or twice a day, for three days before the training procedure started. On the last day of handling, the body weight of the mice was recorded, and food intake was controlled to maintain the body weight at 85% of the initial recorded weight. All procedures followed the European Council Directive (86/609/EEC) and were approved by the Governmental Supervisory Panel on Animal Experiments of Baden Württemberg in Karlsruhe (G-236/20).

### AutoPI task training

Training on the AutoPI task took approximately two months. Mice were first familiarized with the home base for 20 minutes daily for three days. A food reward was delivered every 30 seconds. The lever box was placed inside the home base with the mouse on the fourth day. From this point, food was delivered when the mouse pressed the lever. These daily sessions were terminated after 30 minutes or 100 rewards, whichever came first. The position of the lever inside the home base varied from day to day. After six days of lever training in the home base, the door of the home base was opened, and the lever was placed on the bridge. Over subsequent days, the lever was 2-3 cm further away from the home base until it was located at the center of the arena. Once the lever was at the center of the arena, we trained the mice until they obtained 70 rewards within 30 minutes. Once this criterion was reached, door movements and arena rotations between trials were introduced. During this phase, the mice learned to press the lever independently of its orientation. White noise (65 dB) was also introduced at this point. Then, lever movements were performed every four trials, forcing the mouse to press the lever at different positions and orientations on the arena. The lever only moved between trials.

### AutoPI task

We performed 10-15 behavioral training sessions before the surgical procedure. These sessions included light and dark trials and ended after 60 minutes or when the mouse performed 100 trials, whichever came first. After the surgical procedure, the AutoPI sessions with electrophysiological recordings lasted approximately 90 to 120 minutes. The recording sessions took place in a different room than that in which the AutoPI task training took place.

Every trial on the AutoPi task started with the mouse in the home base and the opening of the door. The mouse then searched for a lever on the arena. Pressing the lever led to the delivery of a food pellet in the food magazine of the home base. The trial ended when the mouse reached the food magazine after pressing the lever or after 5 minutes, whichever came first. The mouse could perform several journeys on the arena during the same trial if the lever had not been pressed during this trial. The mouse was confined to the home base between trials.

Every testing session starts with a consecutive 7 trials under visible light (light trials) followed by an alternating sequence of dark and light trials. All sources of visible light were turned off during dark trials.

The arena was rotated after each trial to one of eight possible angles (multiple of 45°). The lever moved to a new random location on the arena every fourth trial. The combination of arena rotations and lever movement ensured that the lever moved to a new location relative to the home base in most trials. The arena floor and home base were cleaned with ethanol at the end of each recording session. The experimenter monitored the AutoPI session from a different room.

### Surgical procedure

Once the AutoPi task training was completed, the food restriction was paused. The mice were implanted with one H64LP NeuroNexus probe or one or two H10 Cambridge Neurotech probes. Before implantation, the probes were mounted onto custom-made microdrives, allowing independent movement in the dorsoventral axis. Anesthesia was induced with isoflurane (1.5-3%), and the mouse was fixed to the stereotactic frame. Mice were kept under 0.5-1.5% isoflurane during the surgery. The skull was exposed, and two miniature screws were attached to the skull. One screw located above the cerebellum served as a ground/reference electrode. A small craniotomy was performed above the posterior sinus 3.1 mm from the midline. The probes were implanted 0.2 mm anterior of the transverse sinus at an angle of 6-7° in the sagittal plane, with the probe pointing down with a slight angle in the posterior direction. The shanks of the probes were lowered 0.6-0.8 mm into the brain, and the microdrives were fixed to the skull with dental cement. During the first 72 hours after the surgery, the mice received Carprofen (0.1 mg/kg Rymadil, s.c.) every 8 hours. Mice were given at least six days of recovery before food restriction and training resumed.

### Recording sessions

Each recording session started with a 30-minute random foraging trial, during which electrophysiological and positional data were collected as the animal explored a circular arena (diameter = 50 cm). The mouse was moved to a rest box (30 x 30 x 35 cm) for 10 minutes. The AutoPI task was then performed for 90 to 120 minutes. The mouse was then moved back to the rest box for 5 minutes. A second 30-minute random foraging trial was conducted.

After each recording session, the probes were lowered into the brain. For unilateral implantations, the probe was lowered by ∼62.5 µm (Neuronexus Probes) or ∼125 µm (Cambridge Probes). For bilateral implantations, only one probe was lowered on a given day (∼125 µm), alternating between the left and right probes.

On the final day of recording of each mouse, we performed additional recording trials. The lever was placed on the arena for a 15-minute random foraging trial. During this trial, the mouse had no access to the home base of the AutoPI task. The lever’s location on the arena was changed after 15 minutes, and recordings continued for another 15 minutes.

### Histology

At the end of the experiment, the mice were deeply anesthetized using a solution of ketamine (20%; 50 mg/ml) and xylazine (8%; 20 mg/ml) injected intraperitoneally. Transcardial perfusion was performed with 4% paraformaldehyde (PFA). The extracted brains were stored in PFA at 4°C for 24h. The brains were washed with phosphate-buffered saline and cut into 50 µm thick sagittal slices with a vibratome (Leica, Germany). Cresyl violet staining was performed, and stained sections were digitized with a motorized widefield slide scanner (Axio Scan.Z1, Zeiss).

### Electrophysiological recordings, spike extraction, and spike clustering

During electrophysiological recordings, the mouse was connected to a data acquisition system (RHD2000-Series Amplifier Evaluation System, Intan Technologies, analog bandwidth 0.09–7603.77 Hz, sampling rate 20 kHz) via a lightweight cable. The recording hardware was controlled using ktan software (https://github.com/kevin-allen/ktan). Spike extraction and clustering were performed in a semi-automatized manner using Kilosort (https://github.com/jamesjun/Kilosort2). Phy (https://github.com/cortex-lab/phy) was used to refine the spike clusters manually.

Only spike clusters with a distinct waveform, a mean firing rate above 0.1 Hz, and a refractory period ratio smaller than 0.15 were analyzed to ensure high cluster quality. The refractory period ratio was calculated from the spike-time autocorrelation from 0 to 25 ms with a bin width of 0.5 ms. The refractory period ratio was defined as the mean number of spikes in bins 0 to 1.5 ms, divided by the maximum number of spikes in any bin between 5 and 15 ms. We excluded clusters with fewer than 100 spikes in their spike-time autocorrelation (from 0 to 30 ms).

### Recording cable actuator

During recording sessions, the online tracking data were fed to a motorized cable actuator, which ensured that the upper attachment point of the recording cable was kept above the head of the animal. The actuator minimized the interference of the recording cable with the mouse’s behavior. The cable actuator was positioned 153 cm above the arena, with a movement range of 85 x 130 cm in the horizontal plane.

### Position tracking and behavioral analysis

For AutoPI sessions without electrophysiological recordings, we tracked the mouse’s position on the arena with a DeepLabCut model that determined the position of the nose and ears of the mouse. The mid-point between the two ears was used as the animal position. For AutoPI sessions with electrophysiological recordings, an Infrared LED (940 nm) array was attached to the implant and consisted of 3 LEDs on one side and 1 LED on the other side, with a distance of 3 cm between them. The mouse’s position was estimated online using OpenCV functions (for the cable actuator described below). The mouse’s position was recalculated offline using DeepLabCut (https://github.com/DeepLabCut/DeepLabCut) or unetTracker (https://github.com/kevin-allen/unetTracker).

Each AutoPI session was segmented into trials, which started when the door opened and ended when the door closed. The mouse’s path on the arena was divided into search and homing paths from when the mouse pressed the lever. The time when the mouse was located next to the lever was removed from the search and homing path. During the homing path, the mouse was considered to have reached the periphery of the arena when its position was less than 3 cm from the arena’s edge. Trials in which the lever was less than 10 cm from the arena border were excluded from the analysis.

We calculated the length, duration, mean speed, and mean heading direction of the animal during the search and homing paths. Two homing errors were calculated from the homing path—an error at the periphery and a heading error (**Extended Data** Fig. 1a). We calculated the error at the periphery by measuring the angle between the coordinates of the animal at the periphery to the center of the arena and the center of the bridge to the center of the arena. The heading error was defined as the mean heading of the animal during homing until it reached the periphery.

### Identification of grid cells

Firing rate maps were constructed by dividing the area on the circular arena into 3 x 3 cm bins. The number of spikes and the seconds spent in each bin of the maps were calculated, resulting in spike count and occupancy maps. The spike count and occupancy maps were smoothed using a Gaussian kernel (standard deviation = 5 cm). A firing rate map was obtained by dividing the values of the spike count map by those of the occupancy map.

Grid scores were calculated from the spatial autocorrelation of the firing rate map. The location of the six fields surrounding the center of the spatial autocorrelation was obtained to define a circular region in the spatial autocorrelation containing the six fields but excluding the autocorrelation center. The selected region was rotated by 30°, 60°, 90°, 120° and 150°. The rotated regions were correlated with the original, non-rotated region. The grid score was defined 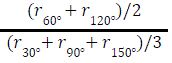, where r is the Pearson correlation coefficient between the non-rotated and rotated regions. The significance levels for grid scores were obtained cell-by-cell by calculating a grid score 500 times after shifting the position data by a random amount (min shift of 20 sec). Grid cells were identified using the first random foraging trial and were defined as neurons with a grid score larger than the 95th percentile of the random grid score distribution.

A grid cell periodicity matrix was generated by performing a correlation between the rotated and non-rotated circular region of the spatial autocorrelation by steps of 2° (**Fig. 1f**). Each row of the periodicity matrix contains the correlation coefficient from one grid cell.

### Reconstruction of movement path from grid cell activity

Recording sessions with at least five grid cells were used for movement path reconstruction. The reconstruction method involved transforming the cartesian coordinate system into a toroidal coordinate system. This transformation depended on the orientation of axes of the grid pattern and their respective periods. We used a multi-step approach to obtain the best estimate for these parameters. We estimated the peak firing rate and grid offset from the firing rate map. The orientation and spacing of the grid cell were obtained from the spatial autocorrelation. These parameters were optimized by fitting the instantaneous firing rate of each grid cell with two grid cell models. The models took the mouse’s trajectory in the open field as input and predicted the firing rate of the neuron for each position data point. The model had three grid axes with their respective direction referred to as θ_0_, θ_1_, θ_2_. The mouse’s position along the three grid axes was calculated (*d*_0_, *d*_1_, *d*_2_). These positions were transformed into angles (*a*_0_, *a*_1_, *a*_2_) using 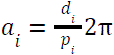, where *p_i_* was the period of each grid axis. The grid pattern of the model had an offset in the three axes (*o*_0_, *o*_1_, *o*_1_). The firing rate of the simulated neuron was obtained with

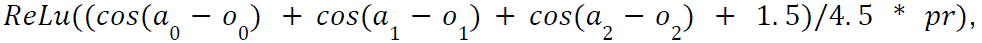

where *p*r was the peak firing rate of the neuron and *ReLu* was a rectified linear unit function. The first model was constrained to have perfect 60° periodicity with axes direction of θ, θ + π/3, θ + 2π/3, and equal axis periods (*p*_0_= *p*_1_= *p*_2_). In the second model, the axis directions were free to deviate from 60° periodicity, and the axis periods could differ. The estimated parameters of the first model were used to initialize the second model. The models were fitted using gradient descent, a mean squared error loss function, and an Adam optimizer. We fitted the two models sequentially to minimize the effect of local minima during gradient descent.

We used the median of θ_0_, θ_1_, *p*_0_, *p*_1_ of the simultaneously recorded grid cells as parameters to transform the cartesian coordinate system into a toroidal coordinate system. The animal’s path on the arena was transformed into a path on the surface of a torus. For each position data point, the position (*d*_0_, *d*_1_) along the first two grid axes was obtained and transformed into angles ʋ_0_, ʋ_1_using 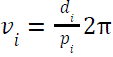. After the transformation, the path of the animal in toroidal space was represented by the angles ʋ_0_, ʋ_1_.

We used a multi-layer long short-term memory (LSTM) recurrent neural network to predict the animal position in toroidal space from the instantaneous firing rate of grid cells. The network had two layers and 256 features in the hidden layers. We used the instantaneous firing rate of grid cells as input (bin size = 20 ms, smoothing kernel standard deviation = 20 ms). The model input was a matrix containing the firing rate of grid cells during the last 400 ms (20 bins), and the model output was a single position on the torus. To deal with the circularity of the coordinates on the torus, the model was trained to predict the cosine and sine of each angle: *cos*(ʋ_0_), *sin*(ʋ_0_), *cos*(ʋ_1_), *sin*(ʋ_1_). The two angles were obtained using *a*r*ctan*2(*sin*(ʋ_0_), *cos*(ʋ_0_)) and *a*r*ctan*2(*sin*(ʋ_1_), *cos*(ʋ_1_)). The model was trained on the first 80 % of the first random foraging trial. The last 20 % were used to test the model using data from the same trial. The model was trained for one epoch, using a batch size of 64, a learning rate of 0.001, a mean squared error loss function, and an Adam optimizer. The predicted *cos*(ʋ_0_), *sin*(ʋ_0_), *cos*(ʋ_1_), *sin*(ʋ_1_) were smooth using a Gaussian kernel (standard deviation = 100 ms).

The sequence of predicted positions in toroidal space was used to calculate the predicted movement path of the mouse in cartesian space. We first calculate the difference Δʋ_0_ and Δʋ_1_ between adjacent ʋ_0_ and ʋ_1_ data points, representing the angular movement along the two dimensions of the torus. Δʋ_0_ and Δʋ_1_ were mapped into movement in cm along the twoaxes using 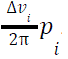. The movement along the two grid axes was then transformed into a movement along the x and y axes of the Cartesian coordinate system.

To assess path integration within the grid module, we calculated the predicted directional error, defined as the angle between the predicted movement vector and the animal’s real movement vector. From the distribution of predicted directional error, we extracted 2 parameters. Directional precision was the mean vector length of the distribution. The circular mean of the distribution represented the orientation of the grid module relative to the first random foraging trial (data used to train the RNN).

### Measurement of grid module anchoring strength

To estimate the location of the lever in toroidal space, we calculated a cartesian vector between the mouse and the lever, transformed this vector into toroidal space, and added the vector to the predicted location of the mouse in toroidal space (**Extended Data** Fig. 5a). To quantify the concentration of the inferred lever positions in toroidal space, we fitted a bivariate von Mises distribution (Kurz & Hanebeck, 2015) to the inferred lever position. The parameters *k*_0_ and *k*_1_ of the bivariate von Mises distribution represent the concentration of the distribution. *k* = 0 for uniform distributions, and larger k values are observed for more concentrated distributions. The mean of the two concentration parameters served as the anchoring strength score. A similar analysis was conducted to assess the anchoring of the grid modules relative to the room reference frame. In this case, the bridge was used as the anchoring point.

### Lever-centered firing rate analysis

We calculated firing rate maps in the lever reference frame. To obtain the position of the animal relative to the lever position, we subtracted the position of the lever from the animal’s position. Firing rate maps were then generated using a procedure similar to that of standard firing rate maps (bin size = 1 x 1 cm), but only data points when the mouse was within a radius of 18 cm from the center of the lever were included. To calculate the map stability during light or dark trials, the trials were divided into two equally sized groups of trials, two firing rate maps were generated, and the Pearson correlation coefficient was calculated between the two firing rate maps.

Directional selectivity around the lever was estimated by calculating the firing rate of a neuron as a function of the direction of the mouse relative to the center of the lever. The direction of the mouse around the lever was defined as the direction of a vector with the origin at the center of the lever and pointing toward the mouse. The firing rate histogram had a bin size of 10°, and only data when the mouse was within a radius of 18 cm from the lever was included. The directional selectivity was defined as the mean vector length of this histogram. To calculate the mean vector length, a 2D vector was obtained for each histogram bin by multiplying the cosine and sine of the angle associated with a bin by the firing rate of the same bin. The 2D vectors were summed and divided by the sum of the firing rate in the histogram. The length of the resultant vector was the mean vector length. To estimate chance levels, directional selectivity was calculated after shifting the position data relative to the spike trains by a random amount (min shift of 20 sec). This procedure was repeated 100 times, and the median was taken.

### Trial matrix of neuron firing rate

We created trial firing rate matrices to analyze the firing behavior of individual neurons on individual trials. Each matrix represented the firing activity of a single neuron. Rows in a matrix represented individual trials. The x-axis represented the animal’s direction relative to the center of the lever (bins of 10°). A smoothing Gaussian kernel (standard deviation = 10°) was applied to each row of the trial matrices.

### Trial matrix of predicted directional error

We constructed trial matrices with the distribution of the predicted directional error for each trial (bins of 10°). The data in the matrices could be limited to the search path, the path at the lever, or the homing path. A smoothing Gaussian kernel (standard deviation = 10°) was applied to each row of the trial matrices. Each row was normalized so that the highest value was 1. A white circle designates the peak predicted directional error within a trial. This peak in the distribution of the predicted directional error was referred to as the trial directional drift.

### Statistics

Wilcoxon signed-rank tests, paired t-tests, and Mann-Whitney rank tests were used to compare two groups. We used Kolmogorov-Smirnov tests to compare the shape of two distributions. For repeated measures, we used the Friedman test to assess differences. Circular-circular correlations were performed as previously described (Jammalamadaka et al., 2001). When appropriate, we used a shuffling procedure to establish significance levels.

## Data availability

The spike trains, position data, and task event logs will be published on the open-access Dryad platform.

## Code availability

The computer code for data analysis will be published on a public GitHub repository.

## Acknowledgments

This work was made possible by the Deutsche Forschungsgemeinschaft via Individual Research Grants (AL 1730/3-1 and AL 1730/4-1 K.A.) and Collaborative Research Centre (SFB-1436, Project-ID 425899996, K.A.). We thank the Lautenschläger Foundation for supporting the Department of Clinical Neurobiology (H.M.).

## Author contributions

K.A. designed the experiments. J.-J.P., B.T., M.N.J., and T.-Y.Y. performed the experiments. K.A., J.-J.P., and B.T. developed and performed the data analysis. K.A., J.-J.P., and B.T. wrote the paper with frequent input from all authors. K.A. and H.M. supervised the project and obtained funding.

## Extended Data

**Extended Data Fig. 1.**
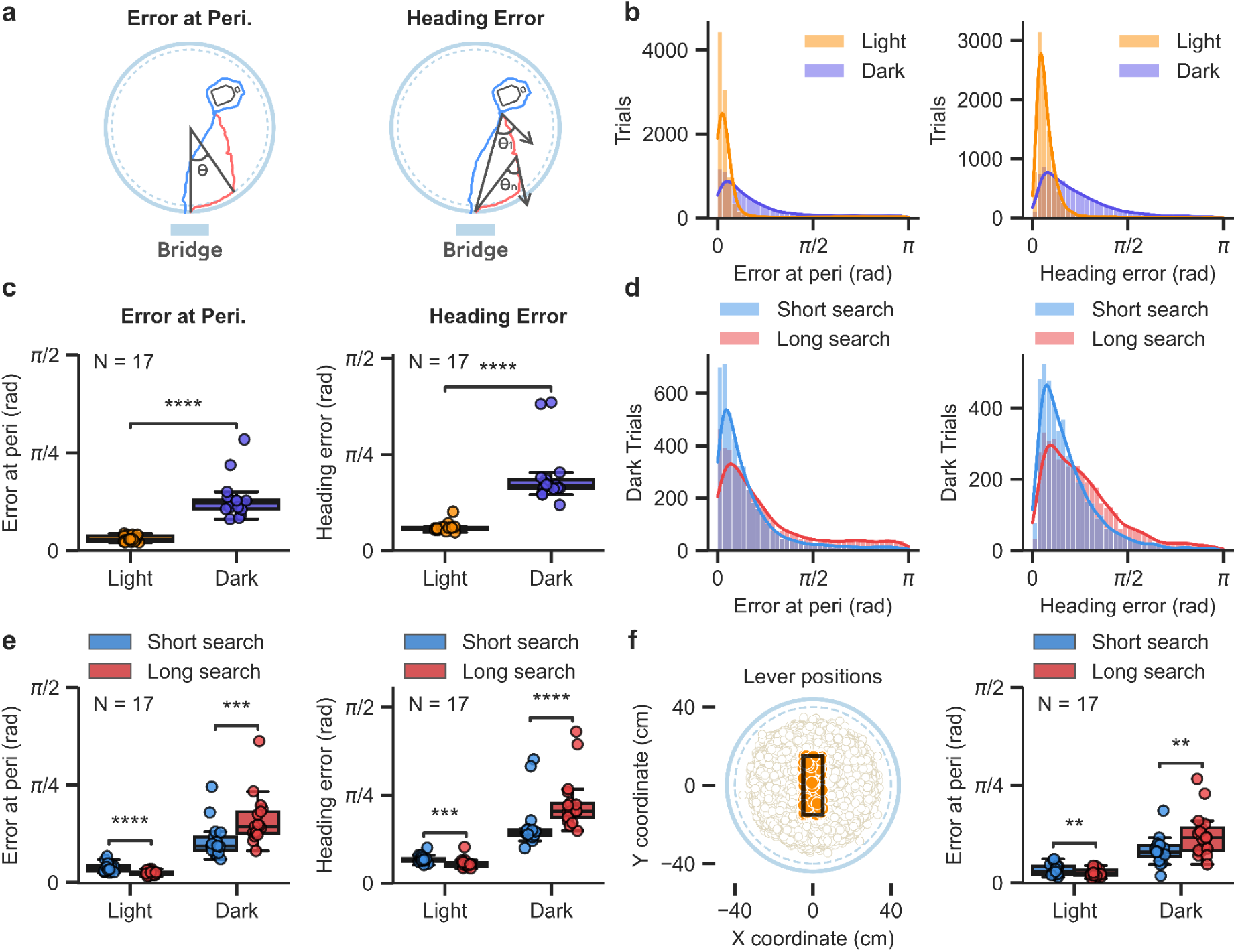
Error at the periphery and heading error of the mouse in different light and search conditions. **a**, Illustration of two measures of homing error: error at the periphery (left) and heading error. Error at the periphery is the angle between the animal’s position when it reaches the periphery of the arena after pressing the lever, the center of the arena, and the center of the bridge. Heading error is calculated along the homing path (at 50 Hz) and is the angle between two vectors with origin at the animal’s position, one pointing in the animal’s direction of movement and the other pointing towards the bridge. The median heading error is calculated for each trial. **b**, Distribution of the error at the periphery (left) and the heading error (right) for homing paths in light and dark trials. **c**, Median error at the periphery (left) or median heading error (right) for each mouse in light and dark trials (N = 17 mice, Wilcoxon signed-rank test, error at the periphery: *P* = 1.526 x 10^-5^, heading error: *P* = 1.526 x 10^-5^). **d**, Distribution of the error at the periphery (left) and heading error (right) for dark trials with short and long search paths. Trials within a session were divided into short and long search paths depending on whether they were below or above the median search path length of the session. **e**, Median error at the periphery (left) or heading error (right) for each mouse during light and dark trials with short and long search paths (N = 17 mice, Wilcoxon signed-rank test, error at the periphery, light: *P* = 1.526 x 10^-5^, dark: *P* = 2.136 x 10^-4^; heading error, light: P = 6.561 x 10^-4^, dark: *P* = 1.526 x 10^-5^). **f**, Effect of search path length during trials in which the lever was located near the center of the arena. We considered only trials in which the lever was located within a 10 x 30 cm rectangle centered on the arena center (left). Error at the periphery for each mouse for long and short search paths in light and dark trials (right, N = 17 mice, Wilcoxon signed-rank test, error at the periphery, light: *P* = 5.569 x 10^-3^, dark: *P* = 6.653 x 10^-3^). ****P < 0.0001, \*\*\**P* < 0.001, ** 0.001 < *P* < 0.01.

**Extended Data Fig. 2.**
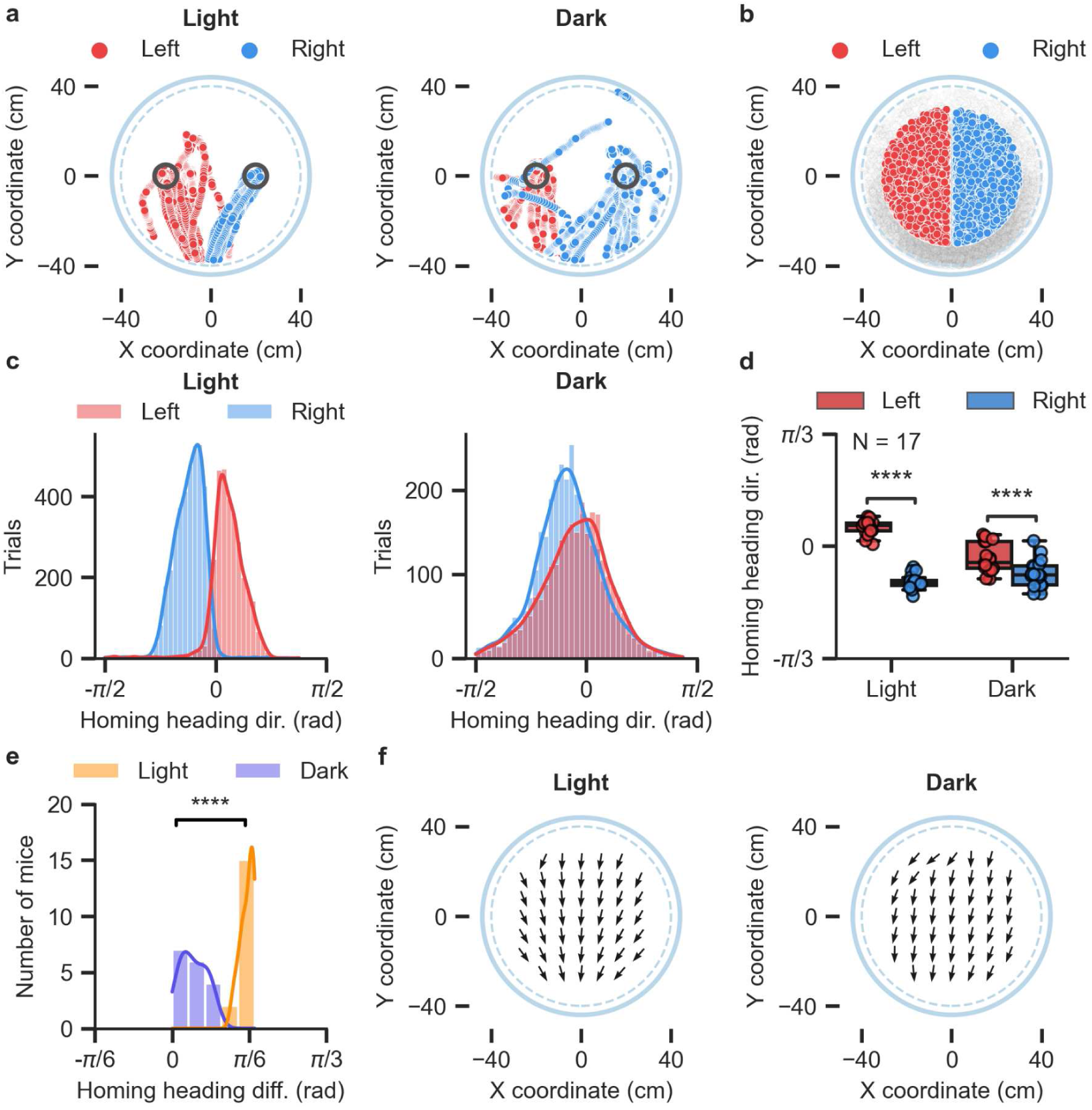
Homing heading direction of the mouse as a function of the animal’s position at the start of the homing path. **a**, Examples of homing paths from one mouse starting at two distinct locations on the arena (two black circles). The homing paths of light and dark trials are shown separately. The arena border is depicted as a gray dashed circle. **b**, Starting position of homing paths grouped based on whether the starting position was on the left or right side of the arena. We excluded starting positions that were more than 30 cm away from the arena center. Starting positions within a 2-cm-wide vertical band at the center of the arena were also excluded. **c**, Distribution of median homing heading direction for homing paths starting on the left or right side of the arena. Data are shown separately for light and dark trials. **d**, Median homing heading direction for each mouse for homing paths starting on the left or right side of the arena (N = 17 mice, Wilcoxon signed-rank test, light trials: *P* = 1.526 x 10^-5^, Dark trials: *P* = 3.052 x 10^-5^). **e**, Difference in median homing direction for homing path starting on the left or right side of the arena. One data point per mouse. The data are shown separately for light and dark trials (N = 17 mice, Wilcoxon signed-rank test, *P* = 1.526 x 10^-5^). **f**, Vector field of the median homing direction of homing paths starting at different positions on the arena. Each arrow represents the median homing direction of homing paths starting at the location of the arrow. The data from all mice are included and shown separately for light and dark trials. ****P < 0.0001.

**Extended Data Fig. 3.**
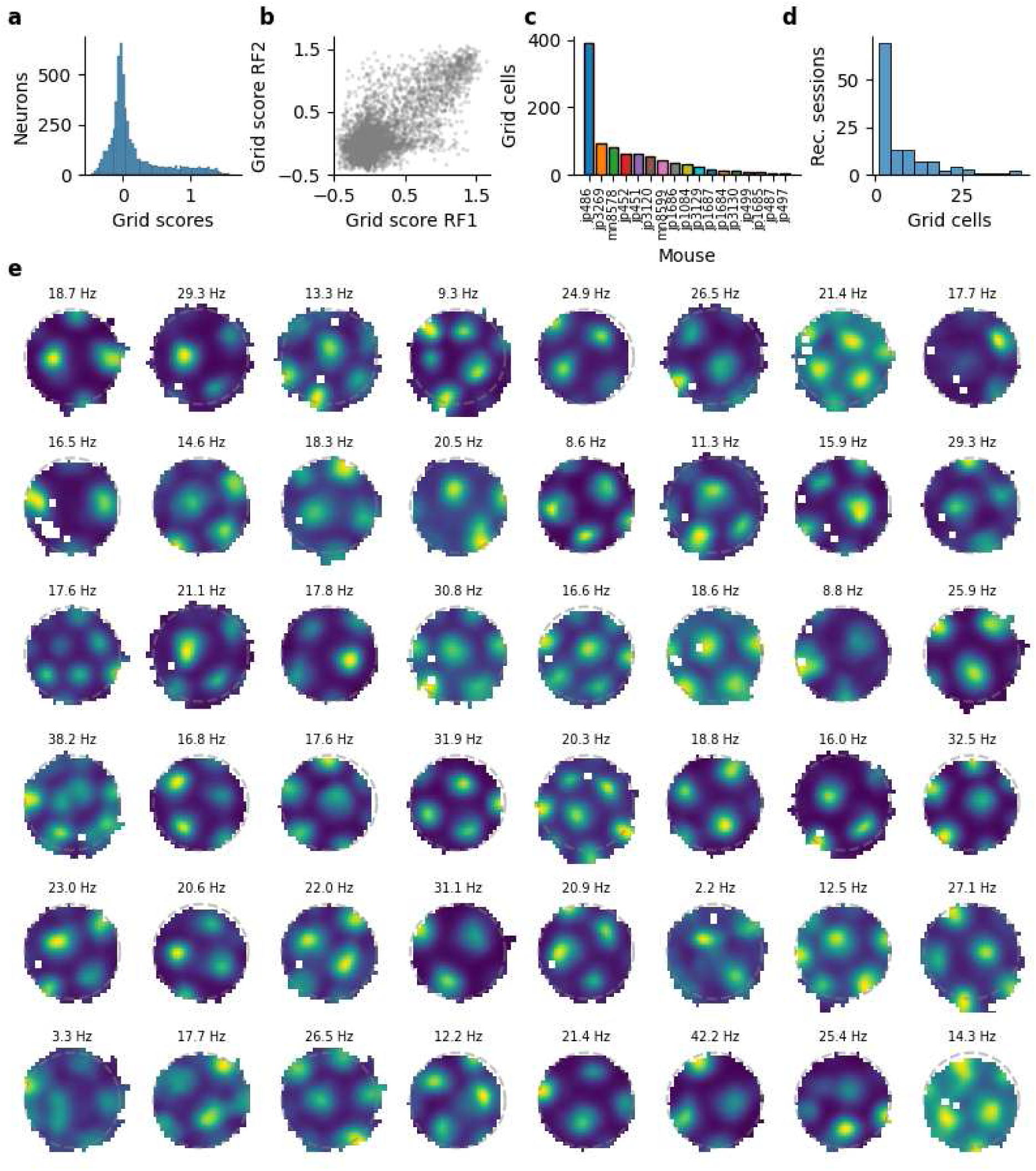
Grid cell identification and number of grid cells per mouse and session. **a**, Distribution of grid scores from all recorded neurons. **b**, Grid scores during the first (RF1) and second (RF2) random foraging trials for all recorded neurons. **c**, Number of grid cells per mouse. **d**, Distribution of the number of grid cells per recording session. **e**, Firing rate map of a random sample of grid cells with a statistically significant grid score during the first and second random foraging trials.

**Extended Data Fig. 4.**
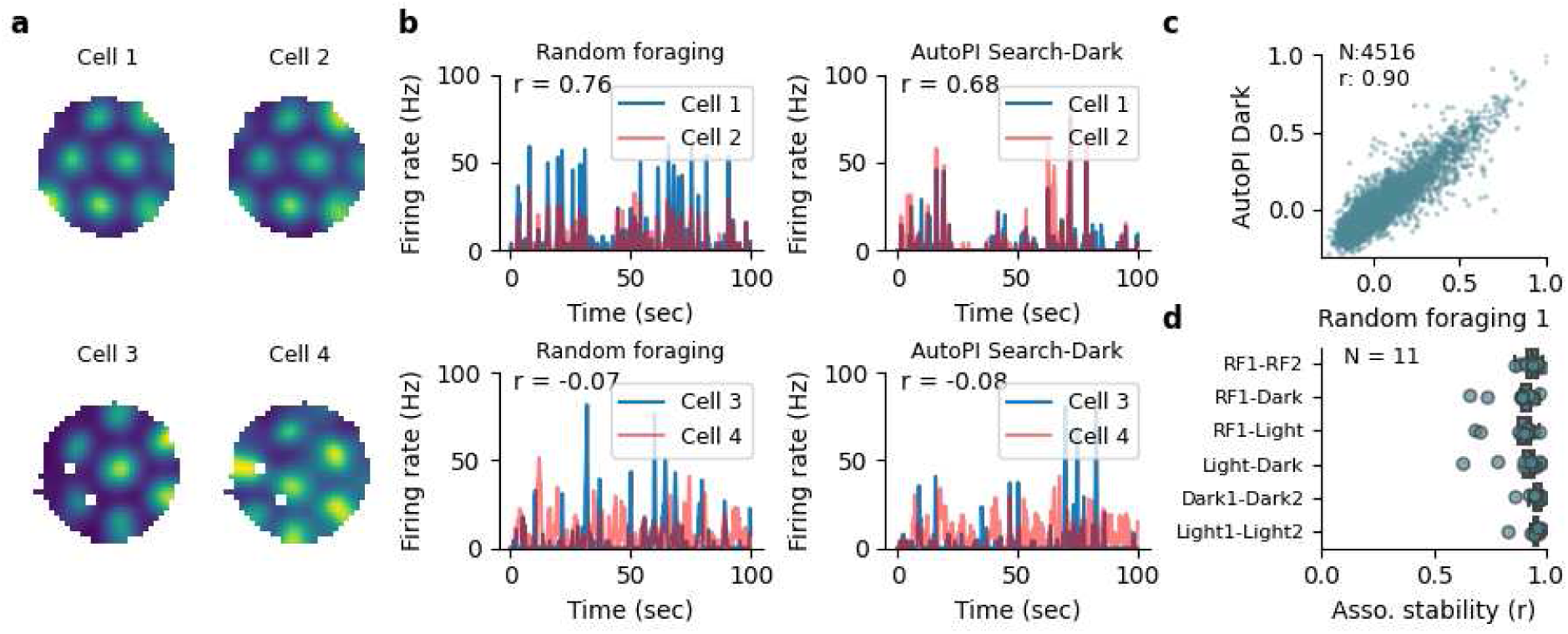
Stability of firing rate associations of grid cells across behavioral tasks. **a**, Firing rate maps of two pairs of grid cells recorded simultaneously. The grid cells on the top row had overlapping firing fields, whereas the ones on the bottom row had non-overlapping firing fields. **b**, Instantaneous firing rate of the grid cell pairs shown in **a** during random foraging and the AutoPI task. The correlation coefficient between the firing rate of the two neurons is shown (r-value). **c**, Firing associations of all recorded grid cell pairs during the first random foraging trial and the AutoPI task (dark trials). Firing associations were strongly correlated across behavioral conditions (N = 4516, r = 0.90, *P* < 1.00 x 10^-16^). **d**, Correlation of firing association for pairs of grid cells across different behavioral conditions (one data point per mouse, 11 mice with at least 10 grid cell pairs).

**Extended Data Fig. 5.**
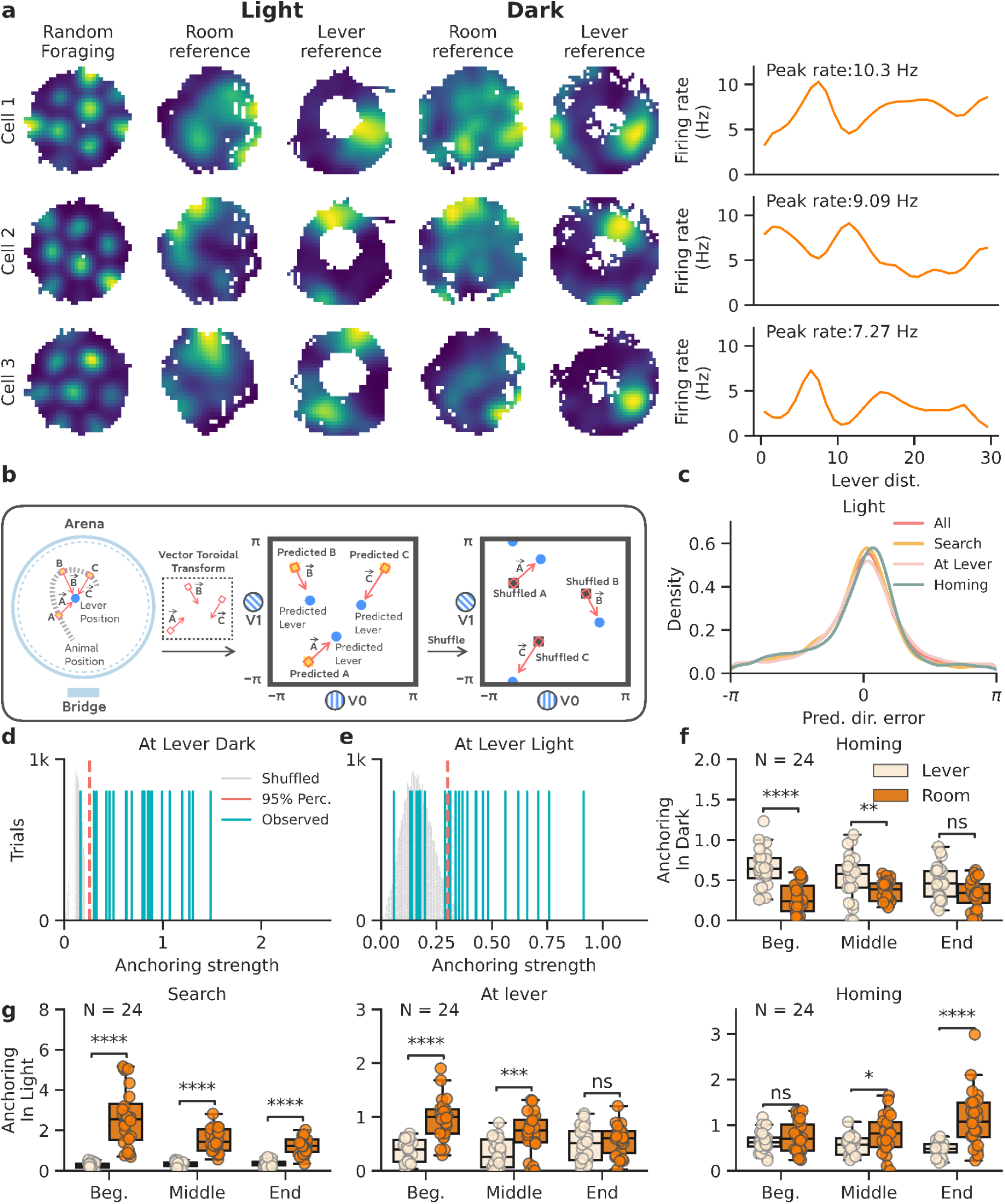
Re-anchoring of grid cell modules to the lever location. **a**, Examples of three grid cells with firing fields near the lever location. Firing rate map of light and dark trials in the arena reference frame and lever reference frame are shown. Firing rate as a function of distance from the lever are also shown. **b,** Graphical abstract of anchoring strength assessment. To measure this, we compute the vector from the animal’s position to the lever and transform this vector into the toroidal space. This vector is added to the decoded animal position as predicted by the model, yielding the predicted lever position for each decoded point. **c,** Probability density map of the angle of rotation between the movement vector prediction of the model against the actual movement vector of the mouse (Directional prediction error). Data is shown for all light trials in four different conditions in the task. **d,** Strength of the grid anchoring to the lever in dark trials. The gray distribution is the anchoring strength expected by chance (shuffling 1000 times). The red line denotes the 95^th^ percentile. Out of 24 sessions, 23 show an above-chance anchoring to the lever. **e,** Strength of the grid anchoring to the lever in light trials. The gray distribution is the anchoring strength expected by chance (shuffling 1000 times). The red line denotes the 95th percentile. Out of 24 sessions, 16 show an above-chance anchoring to the lever. **f,** Anchoring strength to the lever or the room reference during homing in dark trials. The intervals for homing are split evenly into the beginning, middle, and end. **g,** Anchoring strength to the lever or the room reference in light trials. The intervals for search (left) and at lever (middle) and homing (right) are plotted separately and split evenly into the beginning, middle, and end. Data is shown for Search, At Lever, and Homing conditions during the task. ****P < 0.0001, \*\*\**P* < 0.001, ** 0.001 < *P* < 0.01, * 0.01 < *P* < 0.05, ns 0.05 < *P*.

**Extended Data Fig. 6.**
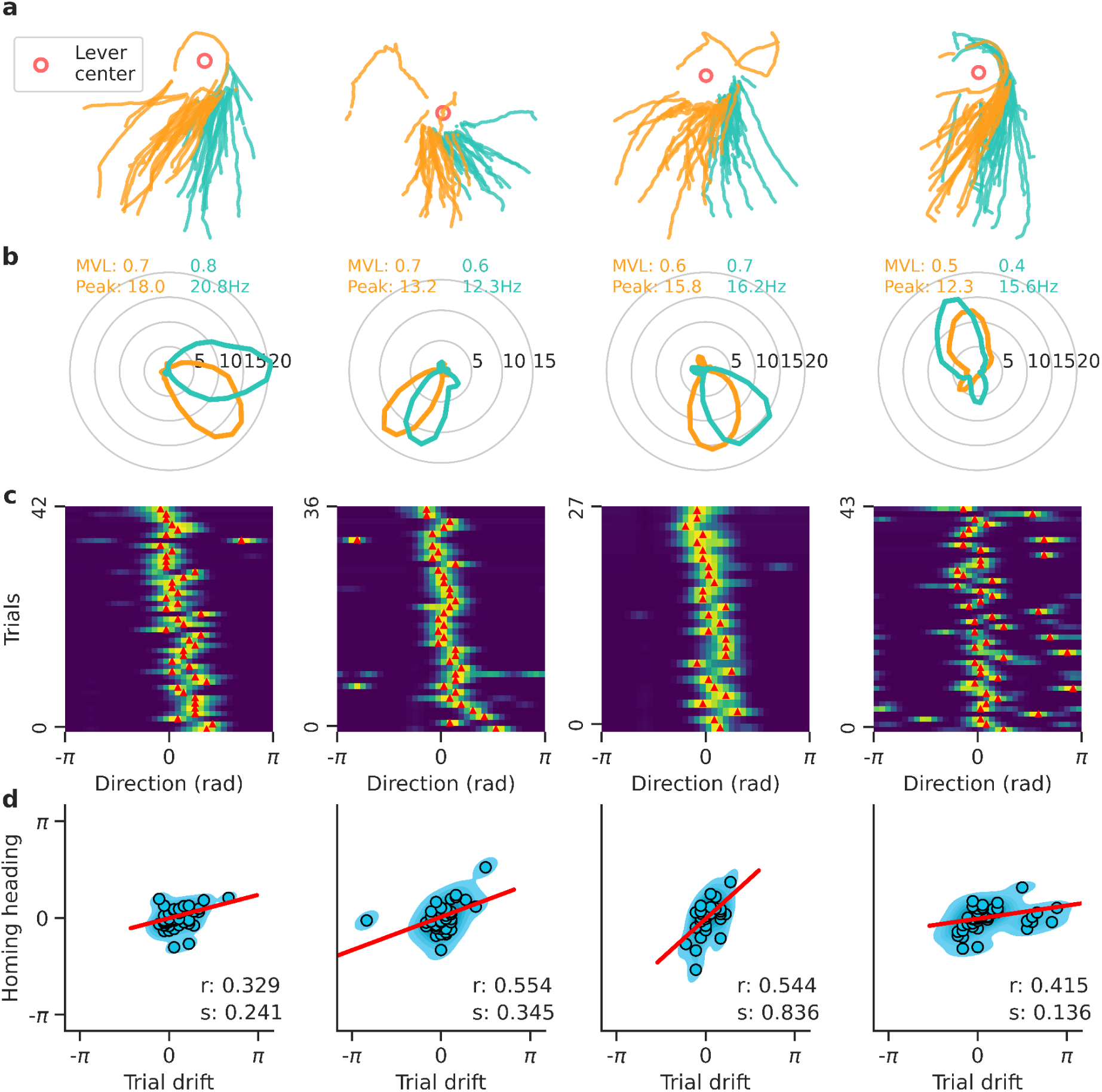
The direction relative to the lever of lever-anchored grid fields predicts homing behavior. Examples of four grid cells with lever-anchored firing fields that predicted homing direction. **a,** trials were divided based on whether the mouse headed left (orange) or right (green) relative to the median homing direction of the session. **b,** Firing rate of four grid cells as a function of the direction of the mouse relative to the lever when the mouse was near the lever. The data are plotted separately for trials in which the animal homed left or right relative to the median homing direction. The mean vector length and peak rate are indicated. **c,** Trial matrix showing the firing rate of the four grid cells as a function of the direction of the mouse relative to the lever. Each row represents one dark trial. The rows were sorted based on the homing direction of the mouse. Red ticks indicate the peak firing direction for each trial. **d,** Trial drift of the neuron against the homing direction of the mouse. Points represent all dark trials of the recording session. A regression line (red), the circular correlation coefficient (r), and the slope of the regression line (s) are shown.

